# Apical-root apoplastic acidification affects cell-wall extensibility in wheat under salinity stress

**DOI:** 10.1101/2020.10.20.347310

**Authors:** Yang Shao, Xiaohui Feng, Hiroki Nakahara, Muhammad Irshad, A. Egrinya Eneji, Yuanrun Zheng, Ping An

## Abstract

Plant salt tolerance is closely associated with a high rate of root growth. Although root growth is governed by cell-wall and apoplastic pH, the relationship between these factors in the root elongation zone under salinity stress remains unclear. Here, we assess apoplastic pH, pH- and expansin-dependent cell-wall extensibility, and expansin expression in the root elongation zone of salt-sensitive (Yongliang-15) and -tolerant (JS-7) cultivars under salinity stress. A six-day 80 mM NaCl treatment significantly reduced apical-root apoplastic pH, from 6.2 to 5.3, in both cultivars. Using a pH-dependent cell-wall extensibility experiment, we found that, under 0 mM NaCl treatment, the optimal pH for cell-wall loosening was 6.0 in the salinity-tolerant cultivar and 4.6 in the salinity-sensitive cultivar. Under 80 mM treatment, a pH of 5.0 mitigated the cell-wall stiffness caused by salinity stress in the salinity-tolerant cultivar, but promoted cell-wall stiffening in the salinity-sensitive cultivar. These changes in pH-dependent cell-wall extensibility are consistent with differences in the root growth of two cultivars under salinity stress. Exogenous expansin application, and expansin expression experiments, we found that salinity stress altered expansin expression, differentially affecting cell-wall extensibility under pH 5.0 and 6.0. *TaEXPA7* and *TaEXPA8* induced cell-wall loosening at pH 5.0, whereas *TaEXPA5* induced cell-wall loosening at pH 6.0. These results elucidate the relationship between expansin and cell-wall extensibility in the root elongation zone, with important implications for enhancing plant growth under salinity stress.

## Introduction

Under abiotic stress, the primary cell wall of plants protects cell integrity and regulates cell expansion and division (Cosgrove, 2018). When plants are exposed to salinity stress, the cell wall, as the outmost layer of cells, is the first line of defense. Salinity stress alters cell-wall structure (Koyro, 1997) and composition (Byrt et al., 2018), factors closely associated with cell-wall extensibility. Salinity stress causes changes in ion homeostasis and transportation across the cell wall. The roots can extrude Na^+^ ions out of cytosol (Munns et al., 2020). Na^+^ extrusion alters the root apoplastic microenvironment, which shifts the apoplastic pH away from the range that favors cell-wall loosening (Byrt et al., 2018).

Studies on changes in apoplastic pH under salinity stress have focused mainly on the leaves. Transient leaf-apoplast alkalization under salinity stress has been reported in the field bean (Felle and Hanstein, 2002), barley (Felle et al., 2005), and maize (Geilfus et al., 2017). Further, an eight-day NaCl treatment induced apoplastic acidification in maize leaves (Zörb et al., 2015). However, there is limited information about the long-term response of root apoplastic pH to salinity stress. Short-term apoplastic alkalization in the root under salinity stress has been reported in *Arabidopsis* (Gao et al., 2004), although that study does not clarify whether the apoplastic alkalization occurred in the root elongation zone or in the mature zone.

Apoplastic pH regulates both cell-wall composition and extensibility. pH-dependent cell-wall extensibility is explained according to the “acid growth theory” (Rayle and Cleland, 1992): apoplastic acidification triggers cell-wall loosening, resulting in cell elongation and expansion. In *Arabidopsis*, apoplastic pH steers root growth (Barbez et al., 2017), promotes cell differentiation (Pacifici et al., 2018), and regulates cell shape (Dang et al., 2020). The optimal pH for cell-wall loosening differs between shoots and roots. In pea (*Pisum sativum*) grown under hydroponic conditions, pH 3.0 buffer induced the maximum apical-root extensibility, compared with pH values of 4.0–8.0 (Tanimoto et al., 2000). In wheat coleoptiles, the optimum pH for cell-wall loosening is 4.0–4.5 (Gao et al., 2008). In maize, root cell-wall extensibility decreased under low water potential (Wu et al., 1996), while low apoplastic pH (pH 4.5) reverses this (Wu and Cosgrove, 2000). However, apoplastic acidification in the root elongation zone does not increase maize growth under salinity stress (Zidan et al., 1990).

Apoplastic pH-dependent cell-wall loosening is facilitated by expansins (Cosgrove, 2005). Cellulose-xyloglucan-cellulose conjunctions form the main load-bearing structures in the cell wall (Park and Cosgrove, 2012); expansin can bind to hydrophilic regions on these conjunctions, unzipping the covalent bonds and loosening the cell walls (Cosgrove, 2018). Expansin genes widely exist in plants. In the wheat genome, the expansin gene superfamily contains over two hundred expansin genes, more than in rice, *Arabidopsis*, and tomato (Han et al., 2019). Many literatures have shown that expansin plays important roles under abiotic stress. In wheat. drought stress makes cell walls more susceptible to exogenous expansin treatment (Zhao et al., 2011). Under low-temperature stress, the expansin gene *TaEXPA8* is highly expressed in a cold-tolerant wheat cultivar (Zhang et al., 2018); in *Arabidopsis*, its overexpression improves cold tolerance (Peng et al., 2019). In wheat under salinity- and PEG-induced stress, expansin gene expression differs between the leaves and roots (Han et al., 2019), and in transgenic tobacco, overexpression of wheat coleoptile expansin genes, such as *TaEXPA2* (Chen et al., 2017) and *TaEXPB23* (Han et al., 2012) enhances root growth and salt tolerance. These studies reveal that, under various types of abiotic stress, expansin acts as a phytohormone and ROS regulator. In contrast, relationships between expansin expression and pH-dependent cell-wall extensibility have not been widely studied.

Therefore, we aimed to clarify how the relationship between apoplastic pH and expansin affects cell-wall extensibility under salinity stress. Using salinity-sensitive and salinity-tolerant wheat cultivars, we examined changes in apical-root apoplastic pH and cell-wall extensibility, using buffers with different pH values and exogenous expansin treatments, and examined expansin gene expression. Understanding the relationship between apoplastic pH and pH-dependent cell-wall extensibility under salt stress is of great importance, first, to elucidate the mechanism of plant growth regulation under salinity stress and, second, to improve the screening or breeding of stress-adapted crop plants.

## Results

### The relationship between root growth and salt tolerance

We treated two wheat cultivars, Yongliang-15 (YL-15; salinity-sensitive) and JS-7 (salinity-tolerant), with NaCl at 0 mM and 100 mM. After 30 d of salinity stress, YL-15 showed worse chlorosis than JS-7(Supplemental Fig. S1). We therefore consider YL-15 to be more sensitive to NaCl than JS-7. In the cultivation experiment using filter-paper, JS-7 showed significantly faster root growth than YL-15, under salinity stress (Supplemental Fig. S1B, C; Fig. 1A, B). After 4 d of 80 mM NaCl treatment, JS-7 seedling root growth recovered and remained relatively high, whereas the root growth rate of YL-15 was low (Fig. 1A). Because the root-growth rates of the two cultivars differed significantly at day 6 of the salinity treatment, we analyzed the apoplastic pH in the roots after 6 d of 80 mM NaCl treatment to investigate the effects of Na^+^ ions on apoplastic pH.

**Figure 1.**
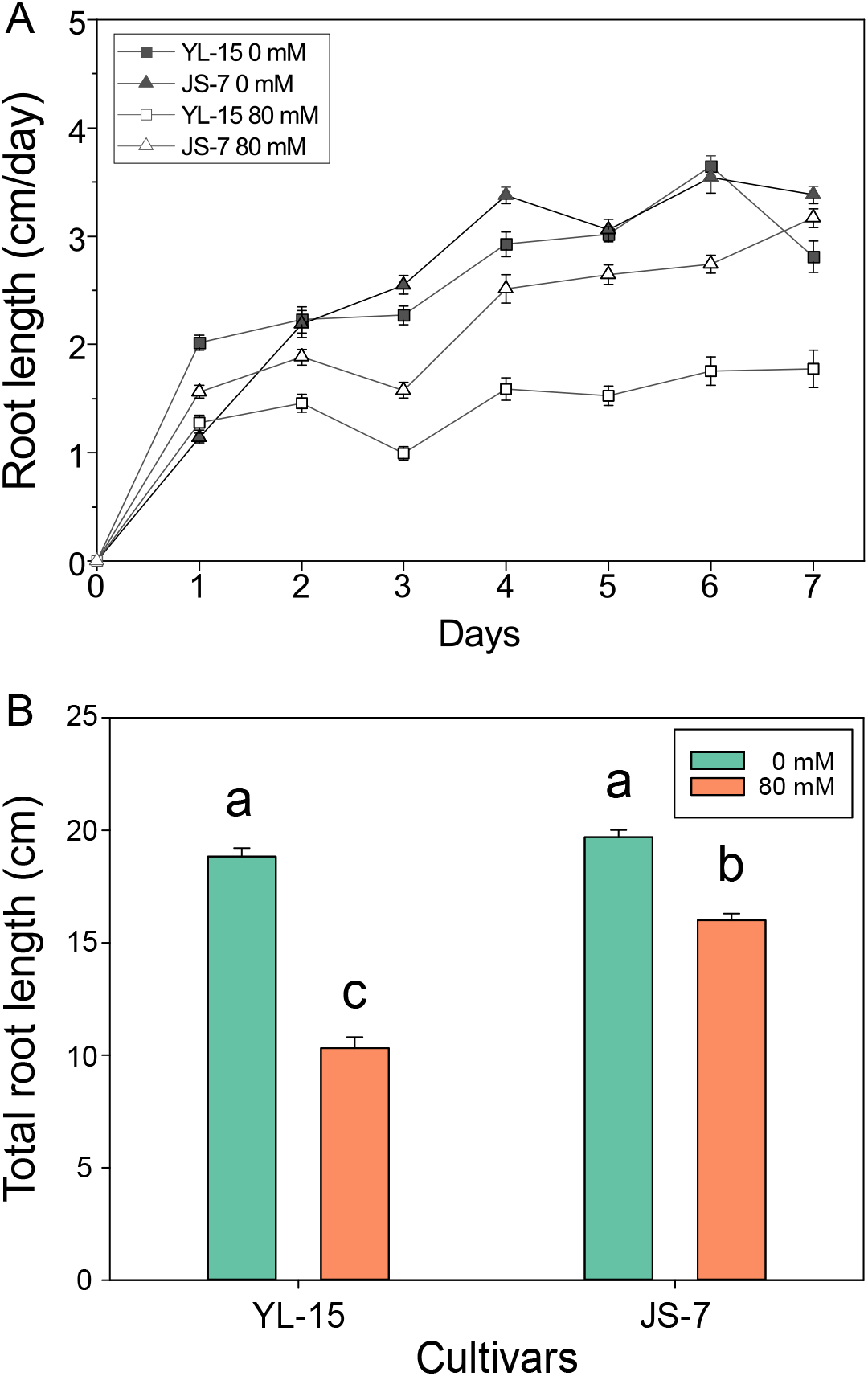
Root-growth rate and total length of the Yongliang-15 (YL-15) and JS-7 wheat cultivars, under the control and salinity-stress treatments. (A) Root-growth rate of YL-15 and JS-7 under seven-day 0 mM and 80 mM NaCl treatments. Growth rate was recorded daily. (B) Total root length of YL-15 and JS-7 under seven-day 0 mM and 80 mM treatments. The data are the mean ± SE (*n*: 37–51).

### Saline stress triggered acidification of apoplastic pH

The 473/405 nm intensity ratio increased with the pH value of the medium (Fig. 2A, B). We used the ratio values to plot a calibration curve ranging from pH 4.0 to 7.0 (Fig. 2C). Moving away from the root tips, the apoplastic pH decreased, and the region 1 mm from the root tips showed the lowest apoplastic pH, in both cultivars and under both of the NaCl treatments (Fig. 2D). Therefore, in this study, we measured pH at 1 mm from the root tip, as representative of the elongation zone. Based on Pritchard et al. (1987), we measured pH at 5 mm from root tip, as representative of the mature zone.

**Figure 2.**
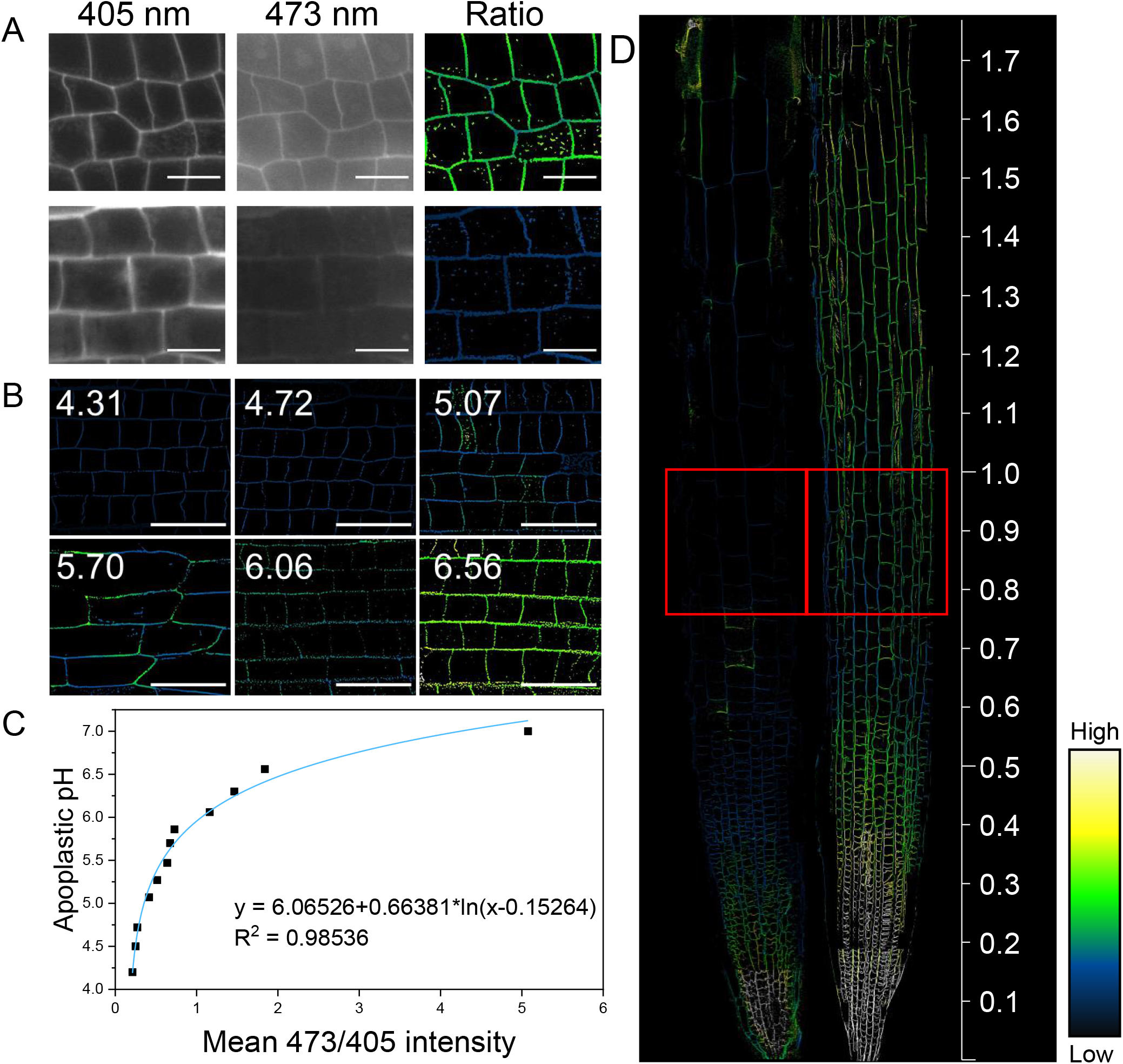
HPTS staining of wheat apical roots, and HPTS calibration. (A) HPTS staining of root cells. (Left) Protonated (acidic) version of HPTS (λ_ex_ 405 nm; λ_em_ 514 nm). (Middle) Deprotonated (basic) version of HPTS (λ_ex_ 473 nm; λ_em_ 514 nm). (Right) Ratiometric image: for each pixel, the 473 intensity is divided by the 405 intensity. (B) HPTS calibration. Apoplastic epidermal root-meristem 473/405 values, of seedlings incubated for 30 min in citrate-phosphate buffer, pH 4.2–7.0. (C) Regression analysis-derived equation enabling calculation of apoplastic pH from the obtained 473/405 values. (D) HPTS-stained root tip of six-day-old seedling under 0 mM and 80 mM NaCl treatments. The color key shows the 475/405 intensity ratio. Scale bars: 20 μm (A) and 50 μm (B).

After irrigating wheat seedlings with 0 mM or 80 mM NaCl supplemented with HPTS (8-hydroxypyrene-1,3,6-trisulfonic acid trisodium salt) for 6 d, we measured apoplastic pH in the root elongation and mature zones. Apoplastic pH was lower in the root elongation zone than in the mature zone in both cultivars under both NaCl treatments (Fig. 3A, B), and apoplastic pH was lower under the 80 mM treatment than in the control, in both zones and cultivars (Fig. 3A, B). Under the 80 mM NaCl treatment, apoplastic pH decreased from ~6.2 to ~5.2 in the elongation zone (Fig. 3B), and from ~6.9 to ~6.1 in the mature zone, in both wheat cultivars (Fig. 3C). In summary, salinity stress induced apoplastic acidification in the elongation and mature zones in both cultivars.

**Figure 3.**
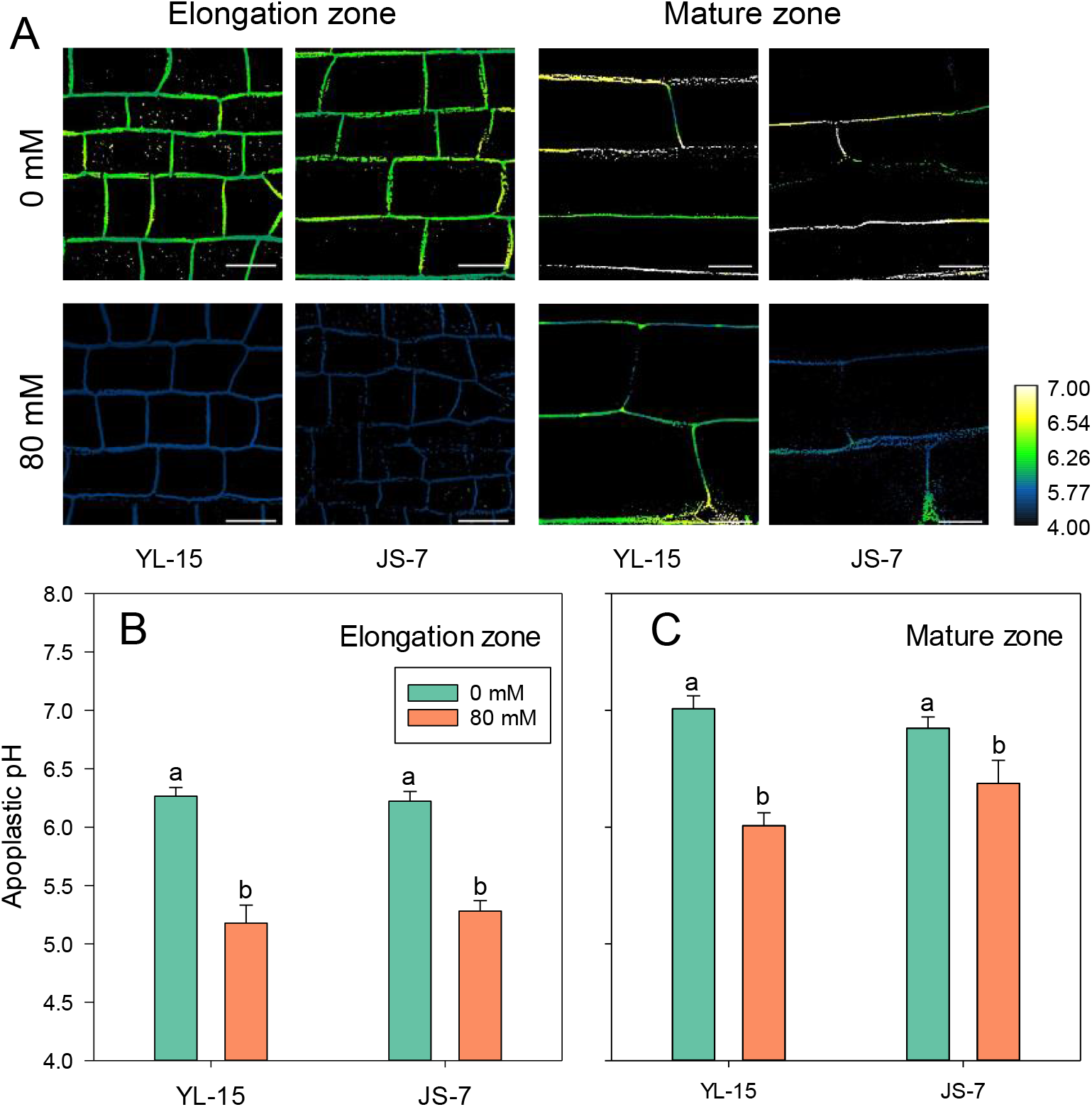
Apoplastic pH in the root elongation and mature zone of Yongliang-15 (YL-15) and JS-7 wheat cultivars, under 0 mM and 80 mM NaCl treatments. (A) HPTS staining of root cells in the elongation (left) and mature (right) zones under 0 mM (top) and 80 (bottom) mM treatments, respectively. The color key indicates pH. Scale bars: 20 μm. (B) Analysis of apoplastic pH in elongation (left) under 0 mM (green bars) and 80 mM (orange bars) NaCl treatments. (C) Analysis of apoplastic pH in the elongation (left) and mature (right) zones under 0 (green bars) and 80 mM (orange bars) NaCl treatments. The data are the mean ± SE (*n*: 6–9 roots per data point). The different lowercase letters above the bars identify groups that differ significantly (*P* < 0.05).

Further, we measured apoplastic pH by soaking the wheat roots in HPTS solution. The HPTS fluorescence signal in the cytosol indicates that salinity stress disrupted cell-membrane integrity (Supplemental Fig. S3A). Apoplastic pH is strongly influenced by the medium’s pH and by hydroponic conditions (Gao et al., 2004). After irrigating the seedlings with the 0 mM or 80 mM NaCl treatment supplemented with HPTS solution (pH 6.5), root elongation-zone apoplastic pH was higher than after the soaking method (Fig. 3, S3B). Further, the HPTS dye was concentrated in the cell walls, under both treatments. Therefore, the cell-wall acidification in the root elongation zone under salinity stress was a direct result of the response of the roots.

### Saline stress altered cell-wall extensibility and pH-dependent cell-wall extensibility in the roots

Byrt et al. (2018) hypothesized that salinity stress alters apoplastic pH, thereby inhibiting cell-wall loosening. To analyze differences in cell-wall extensibility between the two cultivars, we assessed the effects of salinity stress and pH on the cell-wall elasticity modulus (E_0_) and extensibility coefficient (η_N_) of the root tips. Increases in E_0_ and η_N_ are associated with reductions in cell-wall elasticity and creep, respectively (Fig. 4). In both cultivars, E_0_ was significantly higher under the 80 mM NaCl treatment than in the control (Fig. 4A). In general, the increments in E_0_ in YL-15 were greater than those in JS-7, indicating that the root cell walls of YL-15 were stiffer than those of JS-7 under the 80 mM treatment. The differences in E_0_ and η_N_ obtained by varying the buffer pH indicate that, for the 0 mM NaCl treatment, the optimal pH values for cell-wall loosening were 4.6 in the apical roots of YL-15 and 6.0 in segments of JS-7 (Fig. 4A, B); for the 80 mM NaCl treatment, a pH of 4.0–5.0 caused cell-wall stiffening in the apical roots of YL-15, whereas this pH range loosened the cell walls in JS-7 (Fig. 4A, B). In summary, salinity stress stiffened the cell walls and altered the pH-dependent cell-wall extensibility in the apical roots of both cultivars. In contrast, the changes in apoplastic pH inhibited cell-wall loosening in YL-15, but favored it in JS-7.

**Figure 4.**
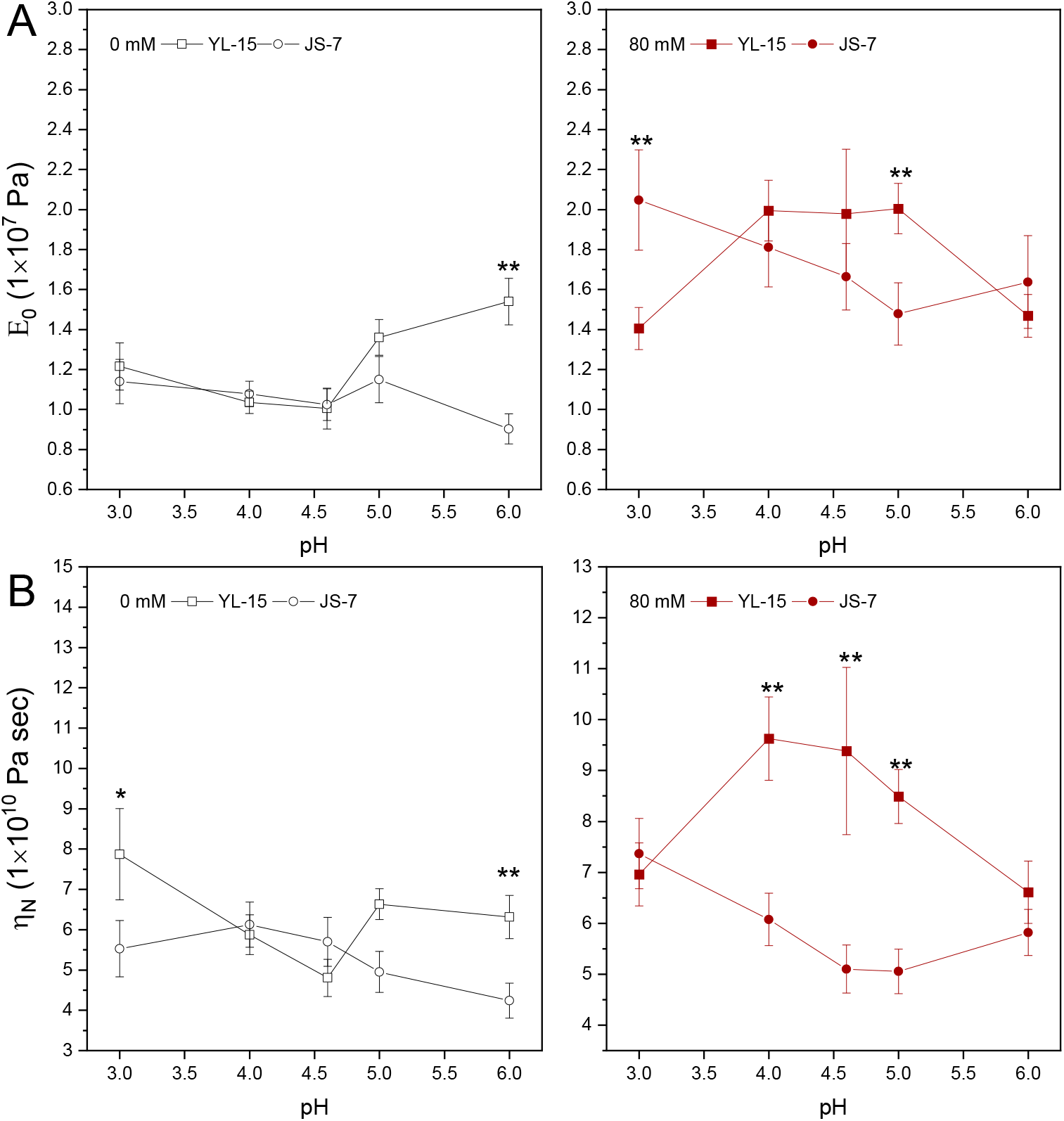
Cell-wall elasticity parameter (E_0_) and creep coefficient (η_N_) in the apical roots of Yongliang-15 (YL-15) and JS-7 wheat cultivars, under the 0 mM and 80 mM NaCl treatments. E_0_ (A) and η_N_ (B) of the cell-wall in the root tips (3–8 mm from root tip) of the two wheat cultivars, at pH 3.0–6.0. The root tips were collected after 10-day 0 mM (left) and 80 (right) mM NaCl treatments. The data are the mean ± SE (*n*: 19–34). Asterisks indicate a significant difference between the wheat cultivars (** *P* < 0.01, *** *P* < 0.001).

### Changes in pH-dependent cell-wall extensibility and expansin expression under salinity stress

We collected expansin-denatured root segments and extracted four sets of expansin samples (from the two cultivars and two salinity treatments). The loosening effects of exogenous expansins on cell walls were assessed under pH 5.0 and 6.0. The expansin extracted from the root of JS-7 grown under the 80 mM NaCl treatment (JS-7 80 mM) induced the lowest E_0_ and η_N_ at pH 5.0, whereas that extracted from YL-15 80 mM induced the highest E_0_ and η_N_ at the same pH (Fig. 5A, B). The apical-root cell walls of two cultivars showed different susceptibilities to the four sets of expansin samples (Fig. 5C, D). The root segments grown under the 80 mM treatment were more susceptible to the expansin extracted from JS-7 80 mM than to that extracted from YL-15 80 mM (Fig. 5C, D). Three-way ANOVA revealed significant effects on cell-wall extensibility of expansin, root segment, and the interaction between expansin and buffer pH, but not of the interaction between root segments and buffer pH (Table 1). The differences in cell-wall extensibility in response to exogenous expansin treatment reveal that expansin determined pH-dependent cell-wall extensibility. When apoplastic pH was at 5.0, expansin from YL-15 80 mM induced cell-wall stiffness, whereas expansin from JS-7 80 mM induced cell-wall loosening.

**Figure 5.**
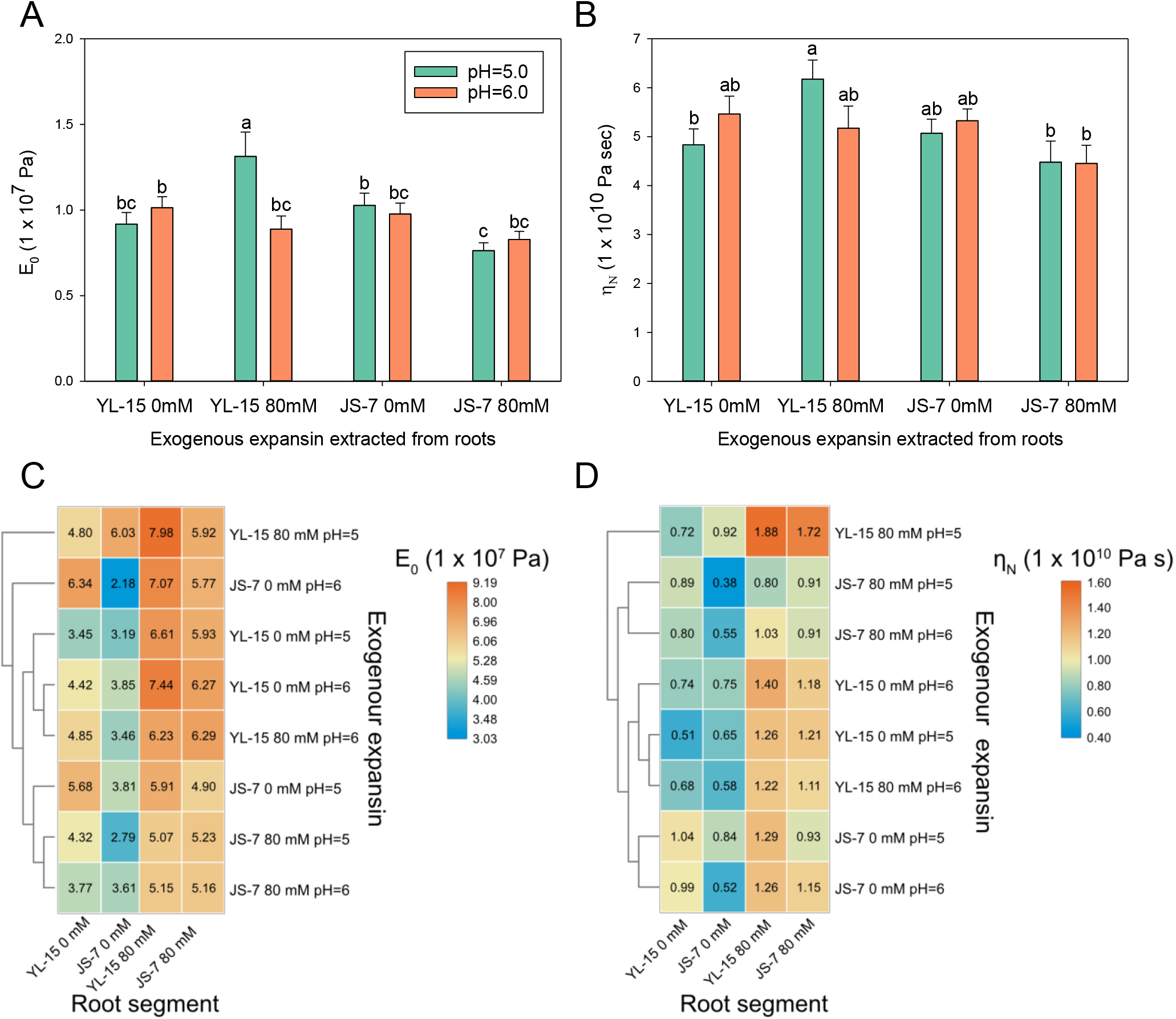
Effects of exogenous expansins on cell-wall elasticity (E_0_) and creep (η_N_) in the root tips of two wheat cultivars, Yongliang-15 (YL-15) and JS-7. E_0_ (A) and η_N_ (B) of apical roots treated with four sets of expansin samples in pH 5.0 or pH 6.0 buffer. The data are the mean ± SE (*n* = 69–79). Expansins were extracted from the Yongliang-15 (YL-15) and JS-7 cultivars under the 0 mM and 80 mM NaCl treatments. The different lowercase letters above the bars identify groups that differ significantly (*P* < 0.05). Heatmap of the E_0_ modulus (C) and η_N_ coefficient (D) of root cell walls treated with the four sets of expansin samples, at pH 5.0 and pH 6.0. Euclidean-distance cluster analysis of the E_0_ and η_N_ data was conducted using TBtools.

**Table 1.** Three-way ANOVA of the effects of root segment, expansin, and buffer pH on the cell-wall elasticity parameter (E_0_) and extensibility coefficient (η_N_). After immersion in exogenous expansin at pH 5.0 or 6.0 for 30 min, the root segments (3–8 mm from the root tip), of plants that had been subjected to 0 mM and 80 mM NaCl treatments, were stretched under 0.05 N tensile force. The exogenous expansins were extracted from the JS-7 and YL-15 wheat cultivars under 0 mM and 80 mM NaCl treatments.

| Dependent Variable: $E_0$ | | | |
| --- | --- | --- | --- |
| Source | df | F-value | Probability |
| Root segment | 3 | 33.500 | 0.000 |
| Expansin | 3 | 7.083 | 0.000 |
| Buffer pH | 1 | 1.772 | 0.184 |
| Root segment * Expansin | 9 | 3.041 | 0.001 |
| Root segment * Buffer pH | 3 | 0.277 | 0.842 |
| Expansin * Buffer pH | 3 | 5.778 | 0.001 |
| Root segment * Expansin * Buffer pH | 9 | 1.253 | 0.260 |

| Dependent Variable: $\eta_N$ | | | |
| --- | --- | --- | --- |
| Source | df | F-value | Probability |
| Root segment | 3 | 25.897 | 0.000 |
| Expansin | 3 | 4.951 | 0.002 |
| Buffer pH | 1 | 0.003 | 0.953 |
| Root segment * Expansin | 9 | 2.089 | 0.029 |
| Root segment * Buffer pH | 3 | 0.976 | 0.404 |
| Expansin * Buffer pH | 3 | 2.196 | 0.088 |
| Root segment * Expansin * Buffer pH | 9 | 1.089 | 0.369 |

We further measured the expansin expression in the two cultivars under salinity stress. Compared with expansin expression in the control, expression was inhibited in the YL-15 80 mM root tips (except for *TaEXPA5* and *TaEXPA9* expression); in contrast, in JS-7 80 mM, *TaEXP5*, *TaEXP7*, and *TaEXP8* showed elevated expression. Under the 80 mM NaCl treatment, the cultivars both showed reduced expression of *TaEXPA3*, *TaEXPA6*, *TaEXPB1*, *TaEXPB7*, and *TaEXPB10*; the reductions did not differ significantly different between the cultivars (Fig. 6). Under salinity stress, elevated *TaEXPA7* and *TaEXPA8* expression mitigated cell-wall stiffness and enhanced root growth in JS-7, the salinity-tolerant cultivar, via apoplastic acidification (Fig. 7). In summary, the roots of JS-7 and YL-15 differentially expressed the expansin genes under salinity stress, which altered the optimal pH for cell-wall loosening.

**Figure 6.**
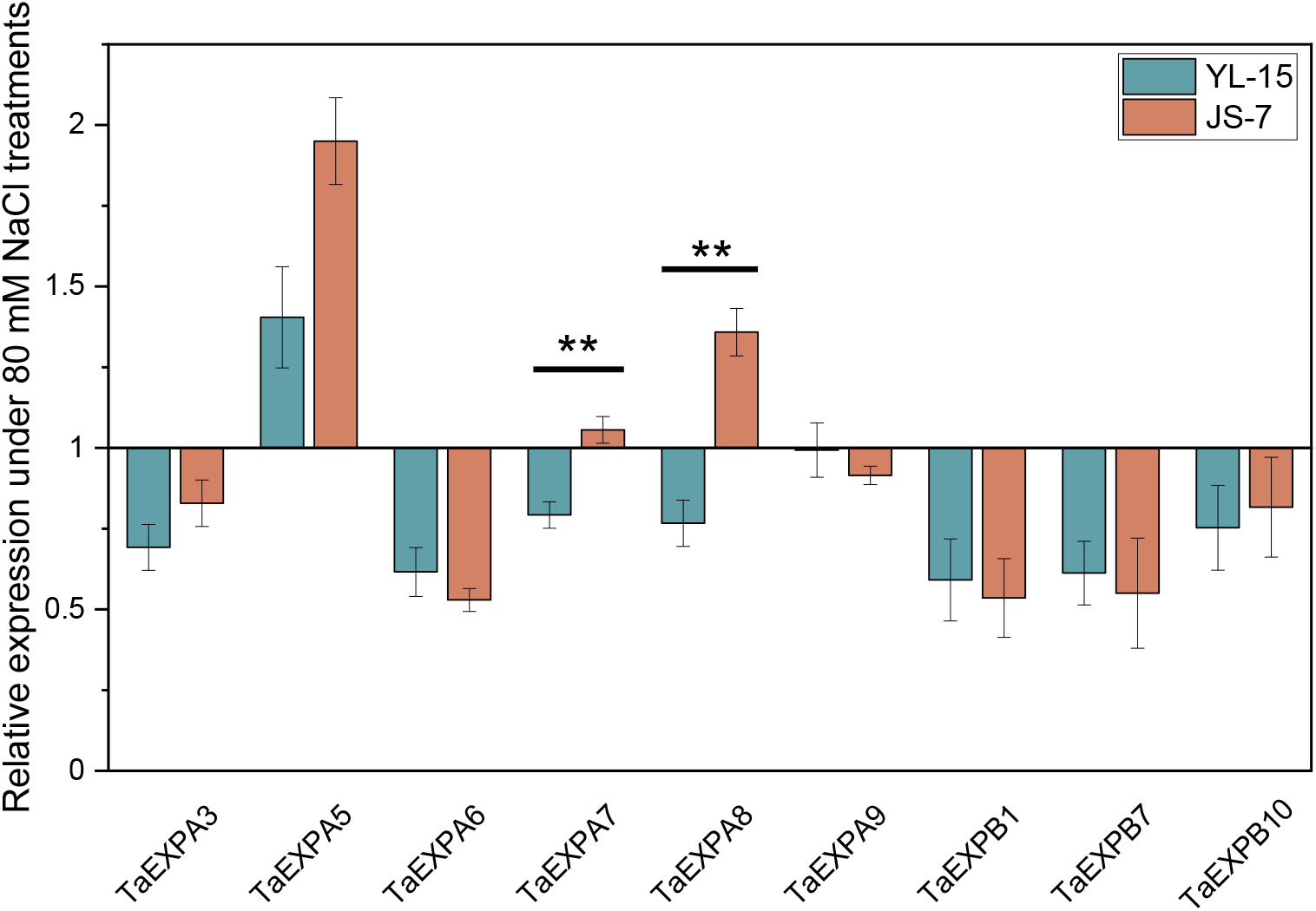
Expression profiling of expansins in wheat roots under 80 mM NaCl stress. The relative expression levels reflect expansin expression under the 80 mM treatment relative to that under 0 mM treatment, in the Yongliang-15 (YL-15) and JS-7 cultivars. Error bars: SEs of three biological replicates. Statistically significant differences between YL-15 and JS-7 were calculated based on Student’s *t*-tests: ** *P* < 0.01

**Figure 7.**
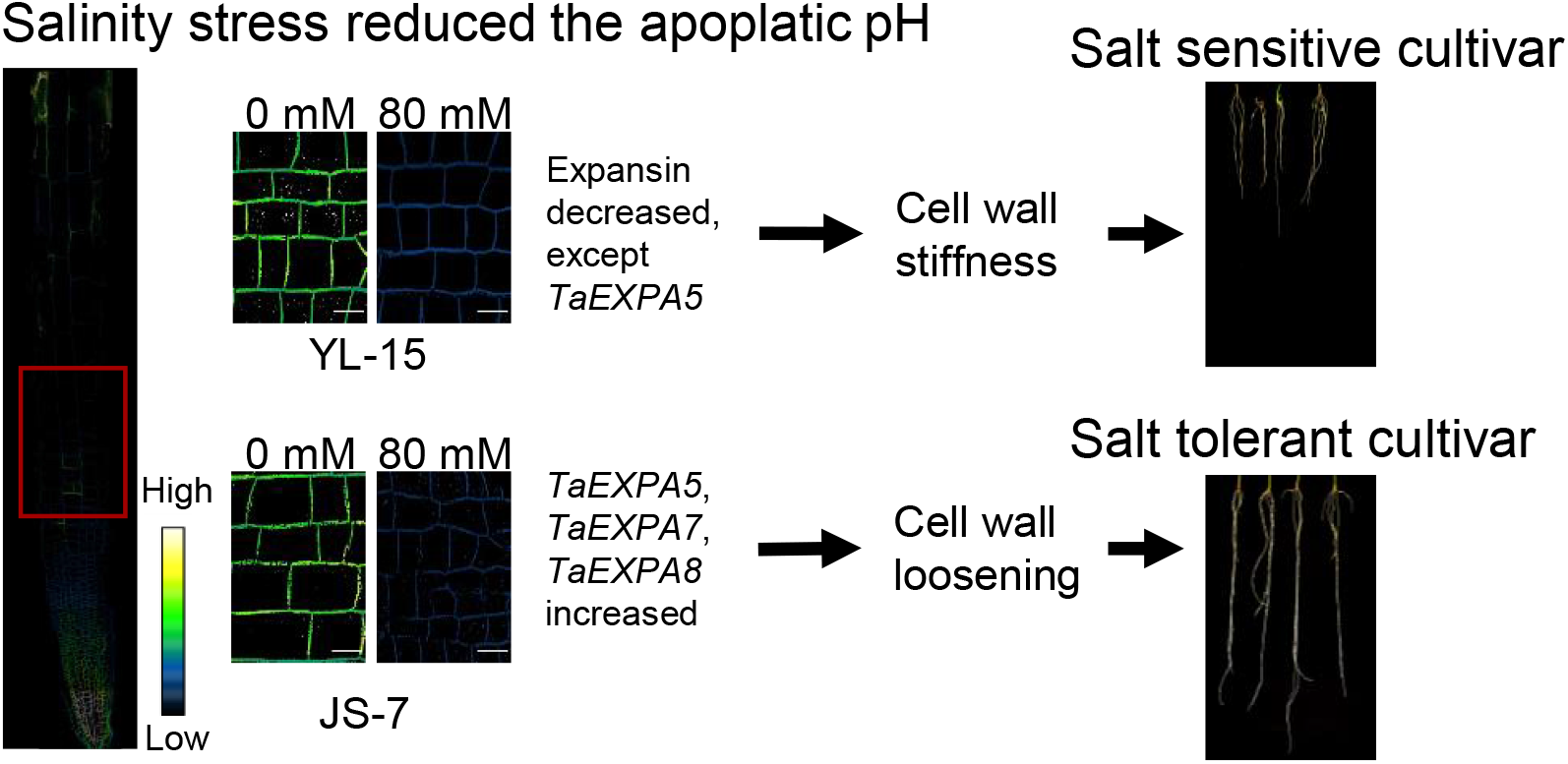
Schematic of the mechanism for mitigating root-growth inhibition under salinity stress in the salt-tolerant wheat cultivar, JS-7. Long-term salinity stress triggers apoplastic acidification in the root elongation zone in both JS-7 and YL-15, the salt-sensitive cultivar. In the salt-tolerant cultivar, Na^+^ triggers elevated *TaEXPA7* and *TaEXPA8* expression, which contributes to cell-wall loosening in the acidified apoplast. This cultivar can thereby maintain a higher root-growth rate under salinity stress than the salt-sensitive cultivar.

## Discussion

### Long-term salt treatment reduced root-tip apoplastic pH

Apoplastic pH steers cell elongation (Barbez et al., 2017), drives cell differentiation (Pacifici et al., 2018), and regulates cell shape (Dang et al., 2020). For both cultivars, we found the apoplastic pH was ~5.2 under 80 mM NaCl condition, whereas the pH was at ~6.2 under 0 mM treatment, indicating that long-term salinity stress significantly acidified the apical-root apoplast. However, previous studies have found that the salinity stress induced transient alkalization in leaf apoplasts—from pH 4.2 to 4.5 in the faba bean (*Vicia faba*; Geilfus and Mühling, 2012) and from pH 4.7 to 5.1 in maize (Geilfus et al., 2015)—and in root apoplasts, from pH 6.4 to 6.8, in *Arabidopsis* (Gao et al., 2004). Leaf apoplastic alkalization plays an important role in stomatal movement (Geilfus et al., 2017). In *Arabidopsis* roots, apoplastic alkalization may be associated with early growth arrest in response to the salinity stress (van Zelm et al., 2020).

The extent of apoplastic acidification caused by long-term salinity stress may be affected by ion-channel functioning and interactions between Na^+^ ions and the cell walls. Ion-channel proteins, such as SOS1 and PM-H^+^-ATPase, together pump more than 95% of Na^+^ ions back into the rhizosphere (Munns et al., 2020). In this Na^+^-extrusion process, PM-H^+^-ATPase is activated to polarize the cell membrane. SOS1 is then activated to pump Na^+^ ions out of cytosol. Further, under salinity stress, Na^+^ ions interact with the polyglucuronic acid (PGA) in cell walls and release the H^+^ from the PGA carboxyl groups (Feng et al., 2018). These findings imply that salinity stress causes apoplastic acidification in roots, especially in root tips, where the pH determines the growth rate of roots.

### Low apoplastic pH reduced cell-wall extensibility in the salt-sensitive cultivar under salinity stress, which severely reduced cell elongation and root growth

The extensibility parameters, E_0_ and η_N,_ were higher in both cultivars under the 80 mM treatment, indicating the salinity stress stiffened the cell walls in both cultivars (Fig. 4 A, B). Thus, the root-growth rates and final lengths of the two cultivars were lower under salinity stress than under the control (Fig. 1 A, B). Under the 80 mM treatment, JS-7 had faster root growth than YL-15, suggesting that JS-7 has superior cell-elongation ability under salinity stress. Interestingly, we found that salinity stress reduced apoplastic pH in the apical roots of both cultivars. However, apoplastic acidification under salinity stress favored cell-wall loosening in JS-7, but had the opposite effect in YL-15. The root growth of YL-15 was severely arrested under salinity stress, more so than that of JS-7 (Fig. 1 A), indicating that changes in the optimal pH for cell-wall loosening are important for root-growth regulation under salinity stress. In summary, salinity stress reduced cell-wall extensibility in both cultivars. However, apoplastic acidification and changes in expansin expression differently regulated cell-wall extensibility in the two cultivars. As a result, under salinity stress, apoplastic acidification further stiffened the root cell walls and slowed root growth in YL-15, whereas, in JS-7, it loosened the root cell walls and mitigated the inhibition of root growth triggered by Na^+^.

### Expansin expression under salinity stress differ between the salinity-sensitive and -tolerant cultivars

We found that the differences in pH-dependent cell-wall extensibility between the sensitive and tolerant cultivars were associated with the differential expression of α-expansins (Fig. 6). Expression levels of *TaEXPA7* and *TaEXPA8* were higher in JS-7 than in YL-15 under salinity stress; in JS-7, expansin caused cell-wall loosening in the acidified apoplast. This suggests that *TaEXPA7* and *TaEXPA8* may contribute to cell-wall loosening under the apoplastic acidification induced by salinity stress. The elevated expression of *TaEXPA5* (Fig. 6) and cell-wall loosening at pH 6.0 (Fig. 5 A, B), in both cultivars under the NaCl treatment, indicate that *TaEXPA5* may contribute to the cell-wall loosening at pH 6.0. Our co-expression analysis showed the *TaEXPA5* regulated the number of roots, whereas *TaEXPA7* and *TaEXPA8* were highly associated with cell lengthening under various stresses (Supplemental Fig. S4). *TaEXPA8* is reported to be closely related to cold tolerance in wheat (Zhang et al., 2018), and the overexpression of *TaEXPA8* improves cold tolerance in transgenic *Arabidopsis* (Peng et al., 2019). These results suggest the importance of *TaEXPA8* in enhancing root growth under various abiotic stresses. A recent study shows that *AtEXPA1* overexpression alters the optimal pH for cell-wall loosening in *Arabidopsis* (Samalova et al., 2020). Therefore, the differential expansin expression between the two wheat cultivars, both under normal conditions and salinity stress, suggests that cell-wall loosening at pH 5.0 and pH 6.0 is caused by different expansin genes: *TaEXPA7* and *TaEXPA8* may induce cell-wall loosening at pH 5.0, whereas *TaEXPA5* may induce it at pH 6.0.

Expansin genes expressed in wheat coleoptiles induce cell-wall loosening under low apoplastic pH (pH of 4.0–4.5; Gao et al., 2008). Further, overexpression of the expansin genes specifically expressed in wheat coleoptiles (such as *TaEXPB23* and *TaEXPA2*) can enhance root growth under salinity stress (Han et al., 2012; Zhao et al., 2012). These results are consistent with our findings. Thus, we conclude that, under the apoplastic acidification caused by salinity stress, the salt-tolerant cultivar elevates its expression of the expansin genes (such as *TaEXPA7* and *TaEXPA8*), which reduces the optimal pH for cell-wall loosening from 6.0 to 5.0. This change in the optimal pH for cell-wall loosening enables the salinity-tolerant cultivar to maintain a higher root growth than the tolerant cultivar.

Further, based on the RNA-seq data from GenBank (accession SRP062745), we found that certain α-expansin, β-expansin, and expansin-like A (*TaEXPLA*) genes are highly expressed only under salinity stress (Supplemental Fig. S5). These stress-specific expansin genes may play a critical role in responses to salinity stress. The α-expansin, β-expansin, and expansin-like A genes related to salinity stress need to be further studied.

### How does the ‘Kelvin-Voigt-Burgers’ model reflect cell-wall extensibility under salinity stress?

Recently, many studies have applied atomic force microscopy (AFM) and Young’s modulus to assess cell-wall extensibility (Feng et al., 2018; Samalova et al., 2020). However, Zhang et al. (2019) report that, although these approaches reveal changes in cell-wall softness, they only partially reveal cell-wall creep and extensibility. AFM assesses cell-wall softness by applying pressure to the cell-wall surface, which does not account for extensibility. In the current study, we found that exogenous expansin application changed root-tip thickness (Supplemental Fig. S6), whereas endogenous expansin had no effect on cell-wall thickness. Samalova et al. (2020) found that expansins were differentially localized in cell walls. Their results indicate that differences in the spatial distribution of expansins regulate longitudinal and horizontal cell-wall loosening, resulting in anisotropic cell growth. Therefore, it is essential to study longitudinal cell-wall loosening in root segments, in order to assess cell extensibility under salinity stress.

Salinity stress induces significant swelling in root tips, increasing the thickness of apical roots (Feng et al., 2018). To avoid this effect when assessing cell-wall extensibility, we used a ‘Kelvin-Voigt-Burgers’ (KVG) model. The KVG model has been used to measure changes in root cell-wall extensibility in wheat roots (Ma et al., 2004) and *Arabidopsis* stems (Shigeyama et al., 2016). The measurements of instant extension and linear extension during constant stretching are used to calculate the E_0_ and η_N_ parameters, respectively. In our study, the root segments were immersed in pH buffers without any force being imposed on them. Therefore, the changes in E_0_ that we observed may relate to cell-wall softness (Fig. 4B, Fig. 5A). The η_N_ parameter, in contrast, reflects the linear extension of cell walls under the constant stretching, which relates to cell-wall creep.

### Salinity stress altered cell-wall susceptibility to exogenous expansins, which was related to cell-wall composition

Drought stress does not change the susceptibility of root cell walls to exogenous expansin in wheat (Zhao et al., 2011). Interestingly, at pH 5.0, the expansin extracted from JS-7 80 mM induced the highest root cell-wall η_N_ extensibility in YL-15 80 mM roots, whereas that from YL-15 0 mM induced the lowest extensibility in YL-15 0 mM roots (Fig. 5 C, D). This indicates that salinity stress alters cell-wall susceptibility to exogenous expansin in YL-15. Changes in cell-wall susceptibility to salinity stress are related to cell-wall structure and composition. Salinity stress alters cell-wall structure, resulting in a stiff and mesh-like network, suggesting an increase in cellulose-xyloglucan conjunctions (Koyro, 1997). The increased xyloglucan then strengthens the linkages between cellulose molecules; this process reduced the growth rate in coffee (*Coffea arabica*) leaf cells under salinity stress (De Lima et al., 2014). Thus, changes in cell-wall susceptibility to expansins may be closely related to changes in cell-wall composition. Further research on changes in cell-wall properties under salinity stress is necessary to elucidate cell-wall susceptibility to expansins.

## Conclusions

Our analyses reveal that apoplastic acidification of apical roots in response to salinity stress stiffens the cell-walls and inhibits root growth in the salt-sensitive wheat cultivar. However, under salinity stress, elevated *TaEXPA7* and *TaEXPA8* expression mitigates cell-wall stiffness and enhances root growth in the salt-tolerant cultivar, via apoplastic acidification.

## Materials and Methods

### Cultivation of wheat seedlings

We used two spring wheat cultivars, Yongliang-15 (YL-15) and JS-7, that differ in salinity tolerance (Supplemental Fig. S1). Seeds of the two cultivars were surface sterilized in 70% ethanol for 5 min, then soaked in distilled water overnight. Twenty seeds were placed in a line on a sheet of filter paper, which was then placed in a 24 × 34 cm Ziploc bag and wetted with 50 ml distilled water. The seeds sprouted in growth chambers (MLR-350 HT, SANYO, Moriguchi, Japan) at 28 °C. Starting two days later, when the roots reached ca. 1.5 cm, 1/12 strength Hoagland solution, containing various treatments, was applied to the roots every 2 d. Light was provided at 2000 lx (16 h light / 8 h dark), and the chamber temperature was constant at 25 °C.

### Apoplastic pH

We used HPTS to investigate apoplastic pH in the elongation and mature zones at a cellular resolution. HPTS, an extracellular pH indicator, has low toxicity because it does not penetrate the cell membrane (Han and Burgess, 2010); it has been used to assess cell-wall apoplastic pH in *Arabidopsis* roots and petals (Barbez et al., 2017; Dang et al., 2020).

For imaging analysis of apoplastic pH in the root elongation zone, 50 ml of 0 mM or 80 mM NaCl solution (1/12 Hoagland solution, pH 6.5), supplemented with 1 mM HPTS, were applied to the two-day-old seedlings. After irrigating the plants with the salt treatments containing HPTS, sin-days-old root of plants were sandwiched between a cover glass and a 35-mm petri dish (with a 20 mm micro-well; Matsunami Glass, Osaka, Japan). Root imaging was performed using a confocal laser scan microscope (FV10-ASW; Olympus, Tokyo, Japan). Fluorescence signals for the protonated HPTS (excitation at 405 nm) and deprotonated HPTS (excitation at 473 nm) were detected using a 60× oil-immersion objective lens. The ratiometric image was obtained by dividing the signal intensity of the 473-nm channel by that of the 405-nm channel, for each pixel. For calibrating the HPTS dye, we stained the wheat roots in medium of a given pH between 4.0 and 7.0, supplemented with 1 mM HPTS, for 30 min. A best-fitting regression method was used to plot the calibration curve. Image analysis was performed using Fiji software (https://fiji.sc/) and a customized macro script (Barbez et al. 2017), with a slight modification (Supplemental Dataset S1). The experiments were performed using at least six biological replicates.

### Expansin extraction

Expansin extraction was performed following Harrison et al. (2001), with a slight modification. Roots (20 g) were homogenized in a blender with liquid nitrogen, and macerated further with 100 ml of extraction buffer (comprising 10 mM 3-[N - morpholino] propanesulphonic acid (MOPS)–NaOH buffer at pH 7.0, 0.5% (w/v) cetyl trimethylammonium bromide, and 30% (w/v) glycerol), until the mixture reached ambient temperature (24 °C). The extraction buffer was then collected and filtered through Miracloth (Merck Millipore, MA, US). Three volumes of precooled acetone were added to the extract, which was then incubated at −20 °C overnight. The mixture was centrifuged at 5 000 ×*g* for 10 min at 4 °C, the supernatant was discarded, and the protein pellet washed once with three volumes of acetone (−20 °C) before freeze-drying. The dried pellets were stored at −20 °C. Protein was assayed using the Bradford method, using a commercial kit (TaKaRa, Shiga, Japan). Before applying to the root segments, each expansin pellet was diluted to 100 μg/ml.

### Root extensibility

As the root extensibility experiments and expansin extraction required many roots (more than 1 000), we extended the experimental period to collect enough samples. Roots tips of ten-day-old seedlings grown in 0 mM or 80 mM NaCl solutions (1/12 Hoagland solution, pH 6.5) were used to measure changes in cell-wall extensibility, at various buffer-pH values and expansin concentrations. Root-tip extensibility was determined following a method developed by Tanimoto et al. (2000). The root extensibility experiments had two parts: Experiment I assessed changes in the cell-wall extensibility of root segments at pH ranging from 3.0 to 6.0; Experiment II assessed the effects of exogenous expansin on cell-wall extensibility using buffer at pH 5.0 or pH 6.0.

In experiment I, root segments of 10 mm from the root tip were prepared by placing them in boiling methanol (80 °C) for 3 min, according to McQueen-Mason et al. (1992). Methanol-boiling treatment denatures the cells but partly preserves expansin activity, whereas water-boiling denatures all expansins (McQueen-Mason et al., 1992). Before the cell-wall extensibility measurement, the methanol-boiled root segments were hydrated with distilled water at ambient temperature (24 °C) for 15 min, then incubated in citrate-phosphate buffer, with pH ranging from 3.0 to 6.0, for more than 30 min under ambient temperature (24 °C). In experiment II, the root tips were boiled in water, to entirely inactive the expansin. To obtain the exogenous expansin for the later experiments, four sets of expansin samples were extracted, one from each cultivar grown under each salinity treatment. After being hydrated with distilled water, the water-boiled root segments were incubated with exogenous expansin at pH 5.0 or 6.0 for more than 30 min.

After being treated with various pH buffers or exogenous expansin concentrations, we measured the extensibility parameters—the elasticity modulus (E_0_) and viscosity coefficient (η_N_)—of the root specimens under a constant tensile force. Before the root specimen was mounted between clips, the diameter of the root at ca. 5 mm from root tip was measured using a microscope (Vitiny UM12, Taiwan, China; Supplemental Fig. S3A). The extensibility of the root region between 3–8 mm from the root tip measured using a creep meter (Yamaden RE2-33005C, Tokyo, Japan). A tensile force of 0.05 N was found to be optimal for obtaining the typical extensibility curve, based on a previous study (Tanimoto et al., 2000) and our preliminary tests. Roots were stretched for 5 min and then released for 5 min. The experiments were performed using at least 19 biological replicates. Details about the root extensibility measurements are reported by Tanimoto et al. (2000).

### Expansin expression and Co-expression analysis

To evaluate changes in expansin expression in response to salinity stress, we extracted the total mRNA from the root tips of YL-15 and JS-7 plants treated with 0 mM or 80 mM NaCl. We then analyzed the transcripts of selected expansin genes that are highly expressed in wheat root tips, which we selected based on a previous study (Lin et al., 2005).

We extracted the mRNA from 60–80 mg of wheat root tips that had been subjected to 0 mM or 80 mM NaCl treatment. mRNA was extracted using a NucleoSpin^®^ RNA kit with rDNase (TaKaRa, Shiga, Japan). cDNA was synthesized using a PrimeScript^®^ RT Reagent Kit (TaKaRa, Shiga, Japan) according to the manufacturer’s instructions. Specific primer sequences for the expansin genes were designed using the Triticeae Multi-omics Center primer server (http://202.194.139.32/PrimerServer; Zhu et al., 2017), and specificity was checked by blasting the sequences against IWGSC RefSeq annotation v1.1 (Appels et al., 2018), shown in (Supplemental Table S1). Actin was used as the reference gene, and we used actin primers from a previous study (Zhu et al., 2016). The qRT-PCR reaction mixture included TB Green^®^ Premix Ex Taq™ II (Takara, Shiga, Japan). The qRT-PCR conditions were as follows: 95 °C for 30 s, followed by 40 cycles of 95 °C for 5 s and 60 °C for 15 s. Three biological replicates were used for each sample. The sequence data were analyzed using the 2^−ΔΔCT^ method, according to the Minimum Information for Publication of Quantitative Real-Time PCR Experiments (MIQE) guidelines (Bustin et al., 2009).

*TaEXPA5-A, -B, -D* (gene ID: TraesCS3A02G165900, TraesCS3B02G199900, TraesCS3D02G175800), *TaEXPA7-A, -B, -D* (gene ID: TraesCS1A02G212000, TraesCS1B02G225700, TraesCS1D02G215100), and *TaEXPA8-A, -B, -D* (gene ID: TraesCS3A02G187600, TraesCS3B02G217000, TraesCS3D02G191300) were examined using the knowledge network generated by KnetMiner (http://knetminer.rothamsted.ac.uk; Hassani-Pak et al., 2020). The expansin network includes both wheat-specific information sources and cross-species information, from *Arabidopsis*. Supplemental Dataset S2 shows how these genes are related to transcription factors and functions. The alluvial diagram, reflecting the relationships between the expansins, their transcription factors, and related functions, was constructed using RAWGraphs (https://rawgraphs.io/; Mauri et al., 2017).

### RNA-seq expression analysis

We used the publicly available RNA-seq data generated from the bread-wheat cultivars Chinese Spring and Qing Mai 6 to study wheat expansin gene expression (GenBank accession SRP062745; Zhang et al., 2016). The expansin expression data were obtained from the Triticeae Multi-omics Center (http://202.194.139.32/expression/index.html). Euclidean-distance cluster analysis of the RNA-seq data was conducted using TBtools (Chen et al., 2020).

### Statistical analysis

Statistical tests were performed using IBM Statistics 21 (SPSS Inc., Chicago, IL, USA). Data-distribution normality was analyzed using the Shapiro-Wilk test. E_0_ and η_N_ were normalized via log-transformation. For pairwise comparisons, statistical differences were detected using a Student’s *t*-test. For comparing apoplastic pH among the zones, cultivars, and treatments, the fluorescence intensity ratio (the intensity at 473 nm divided by the intensity at 405 nm) data were analyzed using a one-way ANOVA and Duncan’s new multiple range test. To detect statistical differences between the groups, in terms of root segments, expansin expression, and pH, we used a three-way ANOVA with Bonferroni post-hoc tests. To compare expansin expression between the cultivars, we used Student’s *t*-tests and Mann-Whitney tests.

## Responsibility

The author responsible for contact and ensuring the distribution of materials integral to the findings presented in this article in accordance with the Journal policy described in the Instructions for Authors (http://www.plantphysiol.org) is Ping An.

## Supplemental data

**Supplemental Figure S1.** Comparison of wheat cultivars Yongliang-15 (YL-15, salinity-sensitive) and JS-7 (salinity-tolerant) under the control (0 mM NaCl) and salinity stress (100 mM for the pot experiment, and 80 mM for the paper experiment).

**Supplemental Figure S2.** Wheat root thickness, measured under the 0 mM and 80 mM NaCl treatments.

**Supplemental Figure S3.** Apoplastic pH in the elongation zone of two wheat cultivars under 0 mM and 80 mM NaCl treatments, after the seedlings were soaked in HPTS solution for 30 min.

**Supplemental Figure S4.** The wheat genes, transcription factors, and functions associated with *TaEXPA5*, *TaEXPA7*, *TaEXPA8*.

**Supplemental Figure S5.** Temporal expression analysis of wheat expansin genes in two bread-wheat cultivars (Chinese spring and Qing Mai 6) roots under NaCl stress.

**Supplemental Table S1.** Primers used for qRT-PCR of wheat expansins.

**Supplemental Dataset S1.** Fiji script for ratiometric image conversion.

**Supplemental Dataset S2.** Transcription factors and functions associated with *TaEXPA5*, *TaEXPA7* and *TaEXPA8*.

## Acknowledgments

The authors are grateful to Mr. Michael Itam of ALRC, Tottori University, Japan, and Dr. Jinghao Zhao of the Rice Research Institute, Sichuan Agricultural University, China, for their helpful advice and discussion.

